# Structural basis of cowpox evasion of NKG2D immunosurveillance

**DOI:** 10.1101/796862

**Authors:** Eric Lazear, Michel M. Sun, Xiaoli Wang, Theresa L. Geurs, Christopher A. Nelson, Jessica A. Campbell, Danna Lippold, Alexander S. Krupnick, Randall S. Davis, Leonidas N. Carayannopoulos, Anthony R. French, Daved H. Fremont

## Abstract

NKG2D is a key component of cytotoxic antitumor and antiviral responses. Multiple viruses evade NKG2D recognition by blocking NKG2D ligand expression on infected cells. In contrast, cowpox virus targets NKG2D directly by encoding a secreted antagonist, Orthopoxvirus MHC Class I-like Protein (OMCP). We have previously reported that OMCP also binds to the orphan receptor FcRL5 on innate B cells. Here, we demonstrate that mammalian-derived, glycosylated OMCP binds NKG2D but not FcRL5. Cowpox viruses either lacking OMCP, or expressing an NKG2D-binding deficient mutant, are significantly attenuated in wild type and FcRL5-deficient mice but not NKG2D-deficient mice, demonstrating that OMCP is critical in subverting NKG2D-mediated immunity *in vivo*. Next we determined the structure of OMCP bound to human NKG2D. Despite a structure similar to that of host NKG2D ligands, OMCP uses a drastically different orientation for NKG2D binding. The re-orientation of OMCP is associated with dramatically higher affinity for human NKG2D and the targeted interface is highly conserved in mammalian NKG2Ds, increasing the zoonotic potential of cowpox virus. We also show that cell surface presented OMCP can trigger NKG2D effector functions equivalently to host NKG2D ligands, demonstrating that NKG2D-mediated signaling requires clustering but is insensitive to binding orientation. Thus, in contrast to TCR/MHC interactions, the docking topology of NKG2D with its ligands does not appear to regulate its activation.

**Author Summary:** Virally infected or tumor-transformed cells display NKG2D ligands (NKG2DLs) on their cell surface, which activates NKG2D-bearing lymphocytes to kill the transformed cell. Pathogens are known to counter this by blocking NKG2DL expression and/or surface display. In contrast, some tumor cells cleave endogenous NKG2DLs creating soluble NKG2D antagonists. Unlike other viral pathogens, cowpox virus uses a strategy analogous to cancer cells by targeting NKG2D directly with a soluble, high affinity NKG2D-antagonist named OMCP. We determined that OMCP’s virulence *in vivo* is attributed to blocking NKG2D-mediated NK cell responses with no apparent effect due to binding to other receptors or cell types. We have also determined the crystal structure of cowpox OMCP bound to human NKG2D, revealing that despite conservation of the ligand scaffolding with host NKG2DLs, the viral protein is engaged with a radically altered orientation compared to all host NKG2DLs. Our structure provides key insight into how OMCP binds with an ∼5,000-fold increased affinity compared to human NKG2DLs and show that the OMCP binding site is exceptionally conserved among primates and rodents, suggesting that the ability of OMCP to recognize this conserved interface contributes to the broad zoonotic potential of cowpox virus. Finally, we show that cell membrane-anchored OMCP can trigger equivalent NKG2D-mediated killing as host NKG2DLs, demonstrating that NKG2D signaling is insensitive to ligand binding orientation.

## Introduction

Intracellular surveillance mediated by MHC class I (MHCI) is a critical host immune function and as such MHCI molecules are frequently targeted for destruction or intracellular retention by viruses [1]. Many herpesviruses encode at least one protein that prevents the cell surface expression of MHCI [1, 2]. However, this immune evasion strategy renders the infected cell susceptible to NK cell-mediated lysis due to loss of inhibitory signals [3]. Viral infection also leads to cell surface display of NKG2D ligands (NKG2DLs) recognized by the activating receptor NKG2D, further predisposing the infected cell towards NK cell-mediated lysis. Therefore, viruses that target MHCI expression often also sabotage NKG2D-mediated cell responses by targeting NKG2DLs on infected cells [4-7].

NKG2DLs are not normally expressed on the cell surface but can be induced by cellular stress [8]. The specific trigger for NKG2DL expression is not known, but NKG2DLs are upregulated in response to several viral infections [9-12]. NKG2DLs comprise a large group of proteins all recognized by NKG2D, despite having low sequence identity. NKG2DLs include the MIC (A and B) and ULBP (1-6) families in humans as well as MULT1 and the RAE-1 (α-ε) and H60 (a-c) families in mice [13]. The redundancy in NKG2DLs is likely due to a combination of tissue specific expression patterns of the ligands and the need to counter viral NKG2D evasion strategies [14]. Many viruses have evolved mechanisms to inhibit the cell surface expression of NKG2DLs as a means of interfering with NKG2D surveillance of viral infection. This strategy is most apparent among β- and γ-herpesviruses, in which four murine cytomegalovirus proteins (m138, m145, m152, m155) [15-18], two human cytomegalovirus proteins (UL16, UL142) [19, 20] and one Kaposi’s sarcoma-associated herpesvirus protein (K5) [21] have been demonstrated to block NKG2DL surface expression. This evasion strategy is also found in RNA viruses, as hepatitis C virus NS3/4a and human immunodeficiency virus Nef proteins also block the expression of a subset of NKG2DLs [22, 23]. Additionally, human cytomegalovirus, herpes simplex virus type 1 and Epstein-Barr virus each also encode at least one miRNA that prevents translation of MICB [24, 25]. Similarly, JCV and BKV polyoma viruses target ULBP3 with miRNAs [26]. However, blocking NKG2DL expression on the infected cell is an imperfect evasion strategy, since no single viral protein or miRNA has been shown to block the expression of all NKG2DLs.

Like several herpesviruses, cowpoxvirus (CPXV) also sabotages MHCI expression. CPXV expresses CPXV012 and CPXV203, two proteins that prevent TAP-mediated peptide transport and MHCI trafficking to the cell surface, respectively [27-34]. Ectromelia virus, a related orthopoxvirus, induces NKG2DL expression, and NKG2D is critical for the control of ectomelia virus pathogensis [35]. Infection with another orthopoxvirus, monkeypox virus, leads to dramatic expansion of NK cells but impaired NK cell function [36]. Together this suggests that CPXV infected cells would be sensitive to NK cell-mediated lysis.

Unlike herpesviruses, CPXV does not target NKG2DLs. Instead this virus targets NKG2D directly by encoding a competitive inhibitor of NKG2DLs, orthopoxvirus MHC class I-like protein (OMCP) [37, 38]. OMCP is a 152 residue protein that is secreted from infected cells and antagonizes the NKG2D-mediated killing of NKG2DL-expressing target cells [37]. OMCP also was reported to bind to B cells via Fc-receptor like protein 5 (FcRL5) and to monocyte/macrophages [39]. OMCP binds to murine NKG2D with an affinity equal or greater than all tested murine NKG2DLs, and to human NKG2D with an affinity ∼5,000-fold higher than human NKG2DLs [37, 38, 40, 41].

Despite their divergence in sequence identity, all known host NKG2DLs share common structural features [42, 43]. NKG2DLs contain an MHCI-like platform domain composed of an eight-stranded beta sheet with two helices [44-48]. The platform domain is subdivided into α1 and α2 domains, with each domain containing four beta strands and an alpha helix. Unlike MHCI, the groove between the helices of the NKG2DL platform domain is closed and therefore NKG2DLs do not bind peptides.

Like host NKG2DLs, OMCP also adopts an MHCI-like platform domain [38]. However, the platform domain of OMCP has been trimmed to have only a six-stranded beta sheet with shorter flanking helices. In this paper, we term the helix of the α1 domain H1 and the discontinuous helix of the α2 domain is termed H2a and H2b. The H2a and H2b helices of OMCP are also rearranged to be flatter against the beta sheet and to be splayed apart from each other. These differences in the OMCP structure were hypothesized to be important for the high affinity binding of OMCP to NKG2D. However, OMCP was still expected to bind to NKG2D in the same orientation as host NKG2DLs, i.e. with the alpha helices oriented diagonally within the symmetric NKG2D binding groove.

Here we report the 2.0 Å-resolution structure of human NKG2D bound to OMCP of the Brighton Red strain of cowpoxvirus. The structure reveals a significant reorientation of OMCP in the NKG2D binding groove relative to host NKG2DLs. The interface of OMCP with NKG2D is highly complementary, buries a significantly larger surface area than host NKG2DLs, and remains continuous across the entire NKG2D binding groove. This novel binding adaptation and high affinity allows OMCP to compete with the high local concentration of membrane-associated host NKG2DLs. We further show that the mechanism of NKG2D antagonism requires OMCP to be secreted, lest it lead to NKG2D signaling. Finally, we show that viruses expressing an NKG2D-binding deficient OMCP mutant are equivalently attenuated to viruses that do not express OMCP using mouse models of cowpox infection. Thus demonstrating that while FcRL5 plays no role in cowpox pathogenesis, OMCP is critical in attenuating NKG2D-mediated immunity.

## Results

### CPXV lacking OMCP is significantly attenuated *in vivo*

While OMCP has been shown to bind to NKG2D and compete with NKG2D ligands *in vitro*, the effect of OMCP on CPXV virulence *in vivo* has not been determined. We generated an OMCP-deficient (ΔV018) CPXV using a transient dominant selection strategy. Multi-step growth curves of the WT CPXV, OMCP-deficient CPXV, and a revertant CPXV demonstrated nearly identical replication growth kinetics (Fig. S1), indicating that OMCP is dispensable for viral growth or replication *in vitro*.

Next we investigated the impact of OMCP during systemic CPXV infection (via the intraperitoneal (i.p.) route) in WT B6 mice. Survival of WT mice infected with WT CPXV demonstrated dose dependence with an LD50 value of 0.91×10^6^ pfu/mouse (Fig. 1A). Comparison of the survival of WT mice infected i.p. with OMCP-deficient CPXV (Fig. 1B) revealed substantially greater survival in the mice infected with OMCP-deficient CPXV than with WT CPX. Indeed, the LD50 value of 1.64×10^6^ pfu/mouse for mice infected with OMCP-deficient CPXV demonstrated that CPXV lacking OMCP is attenuated *in vivo* compared to WT CPXV. Kaplan Meier survival curves of WT mice infected with the OMCP revertant CPXV correlate well with the survival observed following infection with WT CPXV (Fig. 1C), demonstrating that the decreased virulence observed during infection with OMCP-deficient CPXV reflects the absence of OMCP. Together, these results establish for the first time that OMCP facilitates CPXV virulence *in vivo*.

**Figure 1.**
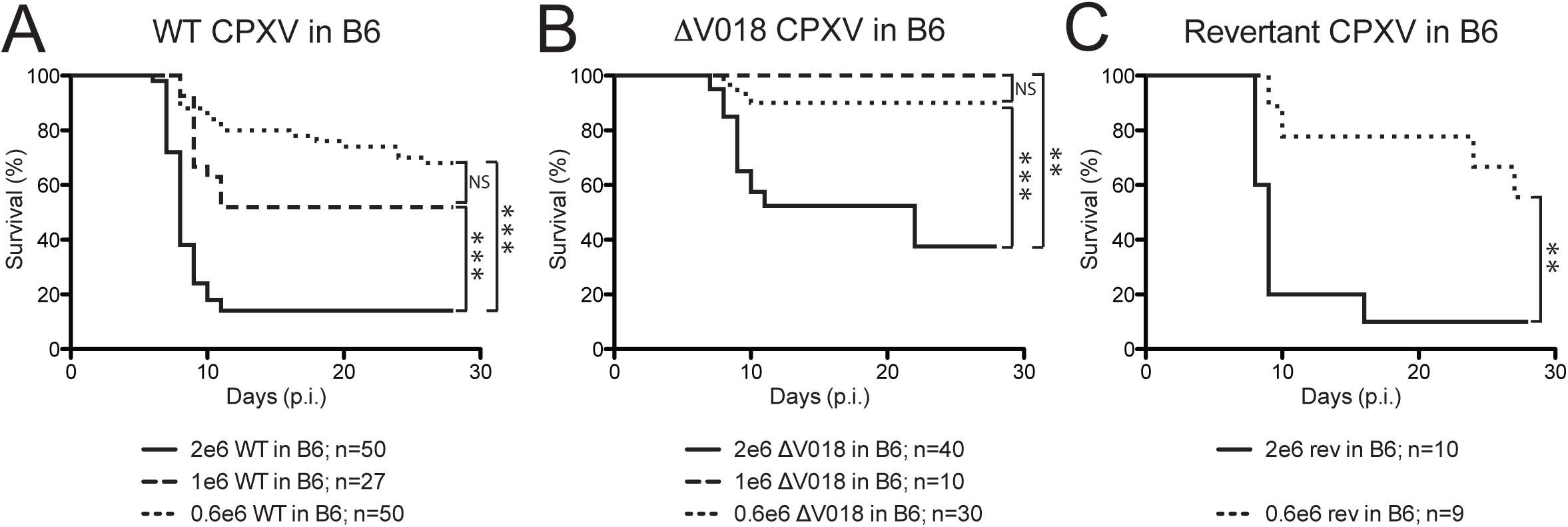
Cowpox virus lacking OMCP is significantly attenuated *in vivo*. WT CPXV **(A)** is more virulent than ΔV018 CPXV **(B)**, as evidenced by increased lethality at all tested doses. The revertant CPXV control **(C)** phenocopies WT CPXV as is expected. 8-12 week old female WT B6 mice were infected with 2×10^6^ (solid line), 1×10^6^ (dashed line), or 0.6×10^6^ (dotted line) of the indicated CPXV strain, and survival was assessed daily for 28 days. Data are aggregated from two-seven independent experiments with 5-10 mice per group.

### Structure determination of OMCP-NKG2D

We had previously solved the structure of OMCP alone and shown that, similar to host NKG2DLs, OMCP adopts an MHCI-like platform domain [38]. Despite the overall similarity of the domain structure of OMCP to host NKG2DLs, OMCP had several notable deviations in the putative NKG2D-binding site that were hypothesized to be important for the high affinity binding of OMCP to NKG2D. To further understand the unusually high affinity of OMCP for NKG2D, we crystallized and solved the structure of OMCP bound to human NKG2D.

Initial crystallization trials with OMCP and NKG2D yielded ∼30 different crystallization conditions. Subsequent data collection and molecular replacement of multiple low-resolution crystal forms all yielded similar partial solutions, with alternating sheets of OMCP-NKG2D complexes separated by undefined density. In the original structure of OMCP alone, the beta sheets packed to form a trimer with the alpha helices oriented away from the center [38]. An identical OMCP trimer formed in the OMCP-NKG2D partial solutions, with NKG2D now bound to the outward facing helices (data not shown). In an attempt to change the lattice packing, we introduced mutations into the beta sheet of OMCP that were designed to break the trimeric interface. These mutations were on the opposite face of OMCP from the NKG2D binding site to avoid disrupting OMCP-NKG2D binding. A mutant form of OMCP (Y23D, F95D) crystallized with NKG2D in a new space group and the crystals diffracted to 2.0 Å (Table 1)(Figure 2A).

**Table 1:** Data collection and refinement statistics^a^.

|  | <u>OMCP<sub>BR</sub>-hNKG2D</u> |
| --- | --- |
| **Data collection** |  |
| Space group | P2<sub>1</sub> |
| Cell dimensions |  |
| a, b, c (Å) | 43.3, 101.1, 91.4 |
| $\alpha\beta\gamma$ (°) | 90.0, 91.6, 90.0 |
| Resolution (Å) | 50-2.0 (2.07-2.00) |
| $R_{\text{sym}}$ | 11.8 (48.5) |
| $I / \sigma$ | 14.5 (3.8) |
| Completeness (%) | 93.5 (91.5) |
| Redundancy | 6.2 (5.3) |
| **Refinement** |  |
| Resolution (Å) | 44-2.0 |
| Total reflections | 309693 |
| Unique reflection | 50139 |
| $R_{\text{work}}$ | 16.6% (21.0%) |
| $R_{\text{free}}$ | 21.4% (29.5%) |
| Wilson B-factor | 21.62 |
| Protein residues | 791 |
| Water molecules | 524 |
| R.M.S. deviations |  |
| Bond lengths (Å) | 0.003 |
| Bond angles (°) | 0.79 |

<sup>a</sup> As defined by PHENIX [75]

**Figure 2.**
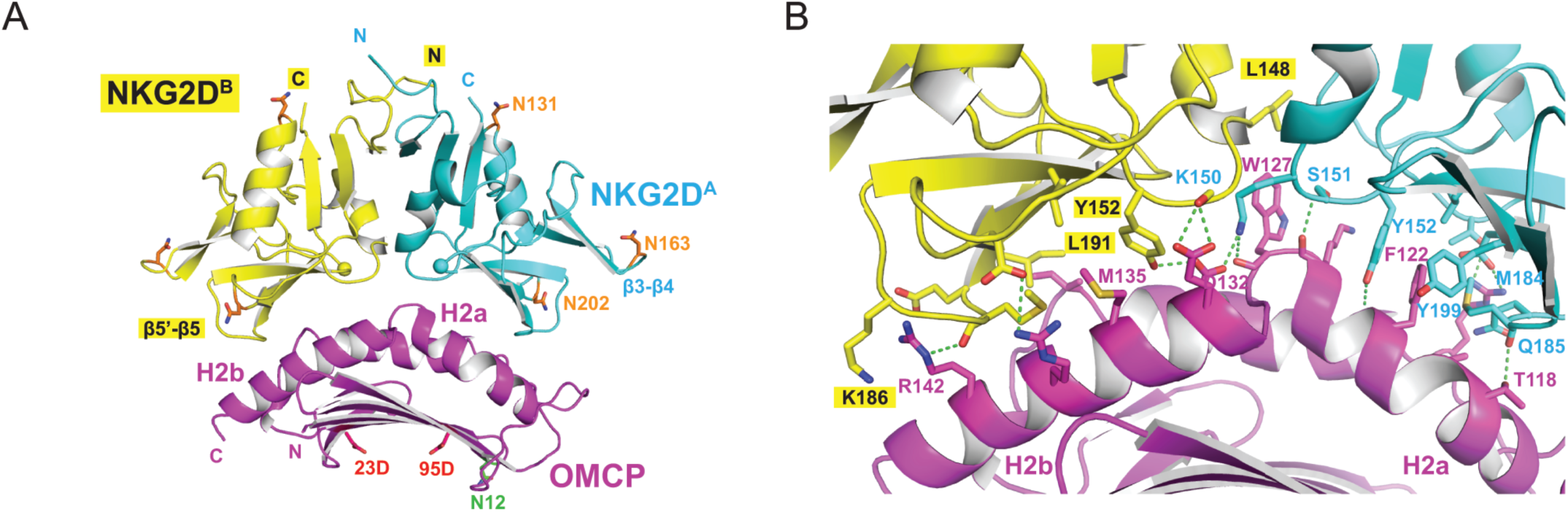
The structure of OMCP in complex with NKG2D. **(A)** OMCP bound to NKG2D. OMCP is colored magenta and the protomers of NKG2D are colored cyan (“A”) and yellow (“B”). NKG2D^A^ makes contacts primarily with the H2a helix and NKG2D^B^ with H2b. Mutations introduced to facilitate alternate crystal packing are shown in red. The S193-S194 bond is shown as a ball on each NKG2D protomer. The asparagines of putative hNKG2D glycosylation sites are shown in orange. The asparagine of the confirmed N-glycan site of OMCP is shown green (data not shown) **(B)** View of the interface between OMCP-NKG2D. The α2 domain of OMCP is shown in the front with the α1 domain behind. OMCP and NKG2D are shown with cartoon representations for the main chain, with the side chains of contact residues shown as sticks. Hydrogen bonds and salt bridges are indicated with green dotted lines.

The electron density map was continuous and unambiguous throughout all chains of the structure, with the exception of Q108 in OMCP. This residue was situated in the center of the largest loop of OMCP and unambiguous density for this residue was also absent from the structure of OMCP alone [38]. The structure of OMCP bound to NKG2D showed no major differences from our previous structure of OMCP alone, with an RMSD for all atoms of 0.8 Å. Likewise, NKG2D was also similar to previous NKG2D structures with RMSDs ranging from 0.5-0.9 Å. The β3-β4 loop of NKG2D is the only region of either OMCP or NKG2D that displayed above-average B factors. This loop is thought to be flexible and has had above average B factors in all previous NKG2D structures [49]. Interestingly, the peptide bond between S193-S194 in our NKG2D structure had a *cis* conformation not described in other NKG2D structures (Figure S2).

### The interface between OMCP and NKG2D

OMCP was hypothesized to bind to the same surface of NKG2D used by host NKG2DLs because (i) OMCP competed with host NKG2DLs for NKG2D and (ii) mutations within the NKG2DL-binding pocket of NKG2D altered OMCP binding affinity [38]. OMCP does bind NKG2D using the same concave binding pocket as host NKG2DLs (Figure 2A). OMCP binds primarily using the discontinuous helices of its α2 domain, H2a and H2b. The position of the H2a and H2b helices is such that every surface exposed side chain of both helices within the binding site directly contacts NKG2D (Figure 2B). Only two contacts are found outside of H2a and H2b, Ile49 and Arg66. Both of these residues are within the α1 domain but lie outside of the H1 helix.

Twelve OMCP residues contact eighteen NKG2D residues to form a mixture of bond types (Table 2). Three residues in each NKG2D half-site are known as core binding residues because they make contacts with all known host NKG2DLs. The core residues of NKG2D subunit A (NKG2D^A^) (Tyr152, Tyr199, Met184) form two hydrogen bonds and make extensive hydrophobic contacts with OMCP residues. The core residues of NKG2D^A^ contact four OMCP residues and the most critical of these residues is Phe122. Phe122 makes multiple hydrophobic contacts with all three NKG2D^A^ core residues, including *pi*-stacking with Tyr152. Phe122 also forms a backbone-to-sidechain hydrogen bond with Tyr152. Interestingly, OMCP is the first NKG2D ligand not to utilize all six NKG2D core-binding residues, with only Met184 and Tyr152 of NKG2D subunit B (NKG2D^B^) contacting OMCP. NKG2D^B^ Met184 and Tyr152 each make a single hydrogen bond and hydrophobic contacts with OMCP residues. Two OMCP residues, Trp127 and Asp132, make contacts with both NKG2D protomers. OMCP Trp127 forms a hydrogen bond to Lys150 of NKG2D^A^ and makes several hydrophobic contacts with Leu148 of NKG2D^B^, Lys150 and Ser151 of NKG2D^A^. OMCP Asp132 forms a hydrogen bond with Tyr152 of NKG2D^B^ and a salt bridge with Lys150 of NKG2D^A^ (Figure 3A).

**Table 2:** Interface contacts between NKG2D and OMCP.

| **NKG2D-A** | **OMCP** | **Bond type** |
| --- | --- | --- |
| Lys150 | Asp132 | Salt bridge |
| Lys150 | Trp127 | H bond |
| Lys150 | Trp127 | Φ (3) |
| Ser151 | Lys126 | H bond |
| Ser151 | Trp127 | Φ (1) |
| Tyr152 | Phe122 | H bond |
| Tyr152 | Phe122 | Φ (9) |
| Tyr152 | Lys126 | Φ (5) |
| Met184 | Thr118 | H bond |
| Met184 | Thr119 | Φ (1) |
| Met184 | Phe122 | Φ (5) |
| Gln185 | Arg66 | Φ (1) |
| Leu191 | Phe122 | Φ (1) |
| Tyr199 | Phe122 | Φ (4) |
| Glu201 | Arg66 | Salt bridge |
| Thr205 | Arg66 | H bond |
| **NKG2D-B** | **OMCP** | **Bond type** |
| Leu148 | Trp127 | Φ (1) |
| Ser151 | Glu131 | H bond |
| Tyr152 | Asp132 | H bond |
| Tyr152 | Glu131 | Φ (3) |
| Tyr152 | Met135 | Φ (5) |
| Ile182 | Ile49 | Φ (2) |
| Glu183 | Arg142 | Salt bridge |
| Met184 | Met135 | Φ (1) |
| Met184 | Arg138 | Φ (2) |
| Met184 | Arg142 | H bond |
| Lys186 | Arg142 | Φ (1) |
| Leu191 | Met135 | Φ (1) |
| Glu201 | Arg138 | Salt bridge |

Hydrogen bonds (H bonds), salt bridges and carbon-to-carbon hydrophobic interactions (Φ) are shown for each contact residue. The number of hydrophobic interactions between contact residues is designated in parenthesis.

**Figure 3.**
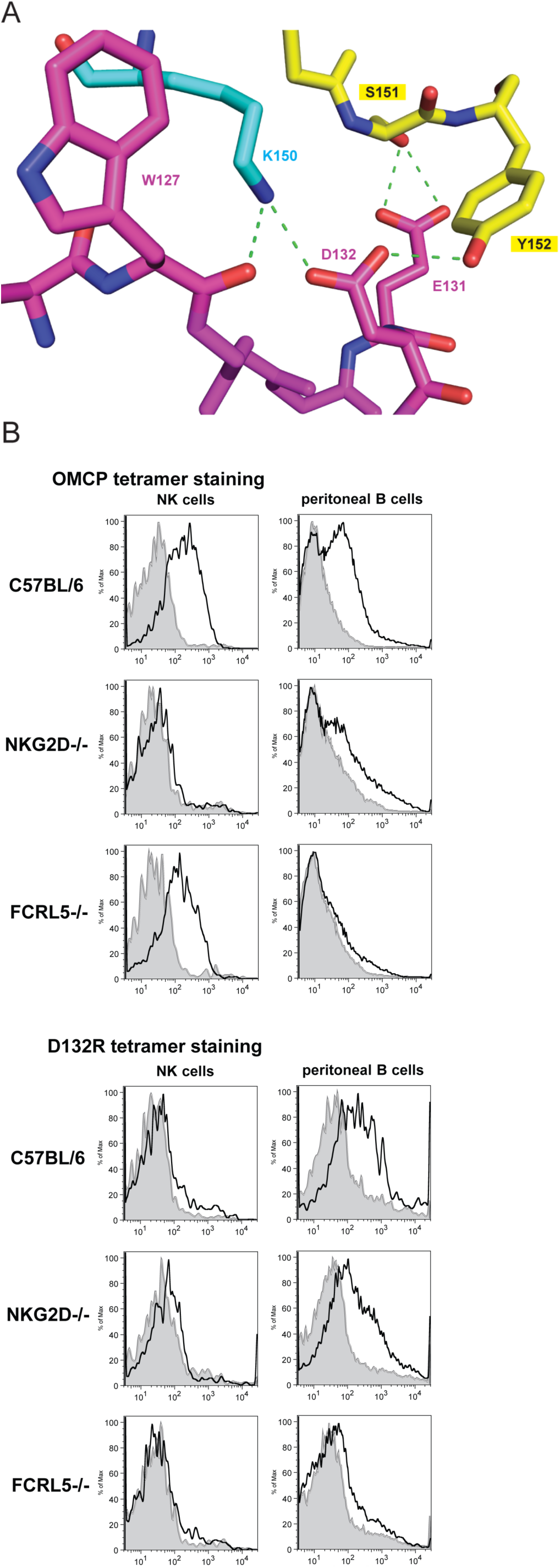
The interface of OMCP and NKG2D. **(A)** The local environment of the OMCP-NKG2D binding interface surrounding the D132R residue. The D132R mutation ablates OMCP-NKG2D binding. **(B)** APC labeled OMCP tetramers (solid line) were used to stain splenic NK cells (NK1.1+, CD3-) and peritoneal B1-a B cells (CD19+, CD5+), Staining with APC labeled WNV DIII tetramer control shown in gray histograms. Representative results from three independent experiments.

Due to the high affinity of the OMCP-NKG2D interaction we harnessed a high throughput *in vitro* selection approach to find NKG2D-binding null mutants (Table S1). The results of the screen identified D132 as an important residue for disrupting NKG2D binding. We then generated OMCP with the D132R mutation in an attempt to completely ablate NKG2D binding. The D132R mutant protein was unable to bind to NKG2D-expressing cells, but retained binding to FcRL5-expressing cells (Figure 3B). This mutation is likely to cause significant steric clashes, as well as disrupting both interactions made by Asp132 to NKG2D^A^ Lys150 and NKG2D^B^ Tyr152 (Figure 3A).

### The role of OMCP receptors in CPXV infection

We generated a recombinant CPXV that expressed the D132R mutant OMCP to isolate the effect of OMCP binding to NKG2D from binding to FcRL5. A multi-step growth curve of (D132R) OMCP CPXV demonstrated identical replication growth kinetics to WT CPXV (Fig. S1). Cowpox and its close relatives variola virus, vaccinia virus, and monekeypox can be spread via contact and aerosol transmission [50]; therefore, we choose to study the role of OMCP during acute intranasal (i.n.) infection. Using our series of WT OMCP expressing CPXV, OMCP-deficient CPXV (ΔV018), and (D132R) OMCP mutant expressing CPXV in combination with WT, NKG2D-deficient, and FcRL5-deficient mice, we were able to delineate the impact of OMCP on NKG2D- and FcRL5-expressing cells during CPXV infection *in vivo*.

Mice were infected i.n. with 2.5×10^4^ pfu/mouse of WT, OMCP-deficient, or D132R CPXV virus. Infection with WT CPXV resulted in similar mortality in WT, NKG2D-deficient, and FcRL5-deficient mice (Fig. 4 solid lines). However, the complete loss of OMCP (Fig. 4, dashed lines) rendered CPXV severely attenuated in WT and FcRL5-deficient mice, while OMCP-deficient CPXV infection in NKG2D-deficient mice remained similarly lethal as WT CPXV. These results illustrated that under normal circumstances (*i.e*., WT CPXV infection in WT mice), OMCP is able to effectively abrogate activation through the NKG2D receptor, rendering WT CPXV as lethal in WT mice as it is in NKG2D-deficient mice. Mice infected with the (D132R) OMCP CPXV (Fig. 4, dotted lines) phenocopied the mortality of mice infected with OMCP-deficient CPXV. The survival observed in the FcRL5-deficient mice following infection with each of the strains of CPXV closely mimicked that seen in WT mice (Figs. 4C vs 4A), indicating that there is no significant role of the FcRL5 receptor on survival following this route of infection. The significant attenuation of OMCP-deficient CPXV and the (D132R) OMCP CPXV infection in WT and FcRL5-deficient mice demonstrated the critical importance of NKG2D blockade during WT CPXV infection, and strongly implicates OMCP blockade of NKG2D as the primary virulence effect.

**Figure 4.**
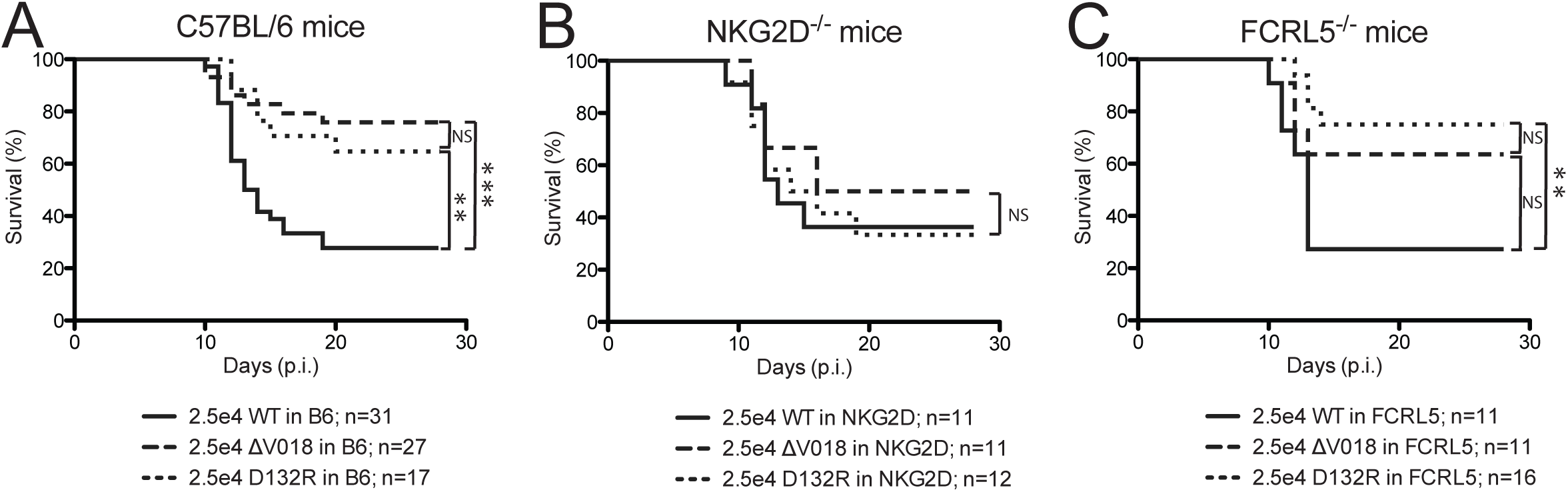
NKG2D mediates all of the virulence of OMCP during intranasal CPXV infection. WT CPXV (solid line), ΔV018 CPXV (dashed line) and D132R OMCP CPXV (dotted line) infected via the intranasal route differentially influence survival in **(A)** WT B6, **(B)** NKG2D-deficient, and **(C)** FcRL5-deficient mice. Kaplan-Meier analysis shows survival of female mice following intranasal infection with 2.5×10^4^ pfu of the designated CPXV strain. The data are aggregated from two to four independent experiments. The Log-rank (Mantel-Cox) test was used in the comparison of all Kaplan Meier survival curves. NS not significant, **p<0.01 ***p<0.001

### Glycosylated OMCP does not bind FcRL5

Compared to previous *in vitro* experiments with bacterial-derived OMCP [37-39], OMCP produced in mammalian cells would be modified by N-linked glycosylation. OMCP is glycosylated at N12, one of its two putative N-glycan sites, when produced in mammalian cells (Fig S3A). The absence of a phenotype for FcRL5 in mice led us to confirm whether mammalian-derived, glycosylated OMCP binds to FcRL5-expressing cells. To test whether glycosylation would affect OMCP receptor binding, we compared binding of bacterially-expressed or mammalian-expressed OMCP to Ba/f3 cells expressing NKG2D or FcRL5 (Fig. 5A). Regardless of source, OMCP bound to NKG2D-expressing cells. In contrast to bacterially-expressed OMCP, mammalian-expressed OMCP did not bind to FcRL5-expressing cells. To confirm whether the difference in FcRL5 binding was due to glycosylation, mammalian-expressed OMCP was treated with EndoF to remove the N-linked glycan. The deglycosylated protein gained the ability to interact with FcRL5. Next we tested the binding of mammalian-expressed OMCP to primary human PBMCs (Fig 5B). As a positive control for NKG2D binding, we included the human NKG2DL ULBP3. WT OMCP and ULBP3 stained both NK and a subset of T cells, but not B cells, consistent with NKG2D expression on these cells. In contrast, (D132R) OMCP failed to significantly stain NK and T cells, supporting that WT OMCP cell staining is due to binding NKG2D. Together the *in vivo* virulence (Fig 4) and cell binding experiments (Fig 5) demonstrate that mammalian-derived OMCP functions only as a high affinity NKG2D antagonist and is not a bona fide FcRL5 ligand. Importantly, OMCP blockade of NKG2D plays a significant role in pathogenesis in mice, and potentially an even larger role with higher affinity NKG2D orthologs, like human NKG2D.

**Figure 5.**
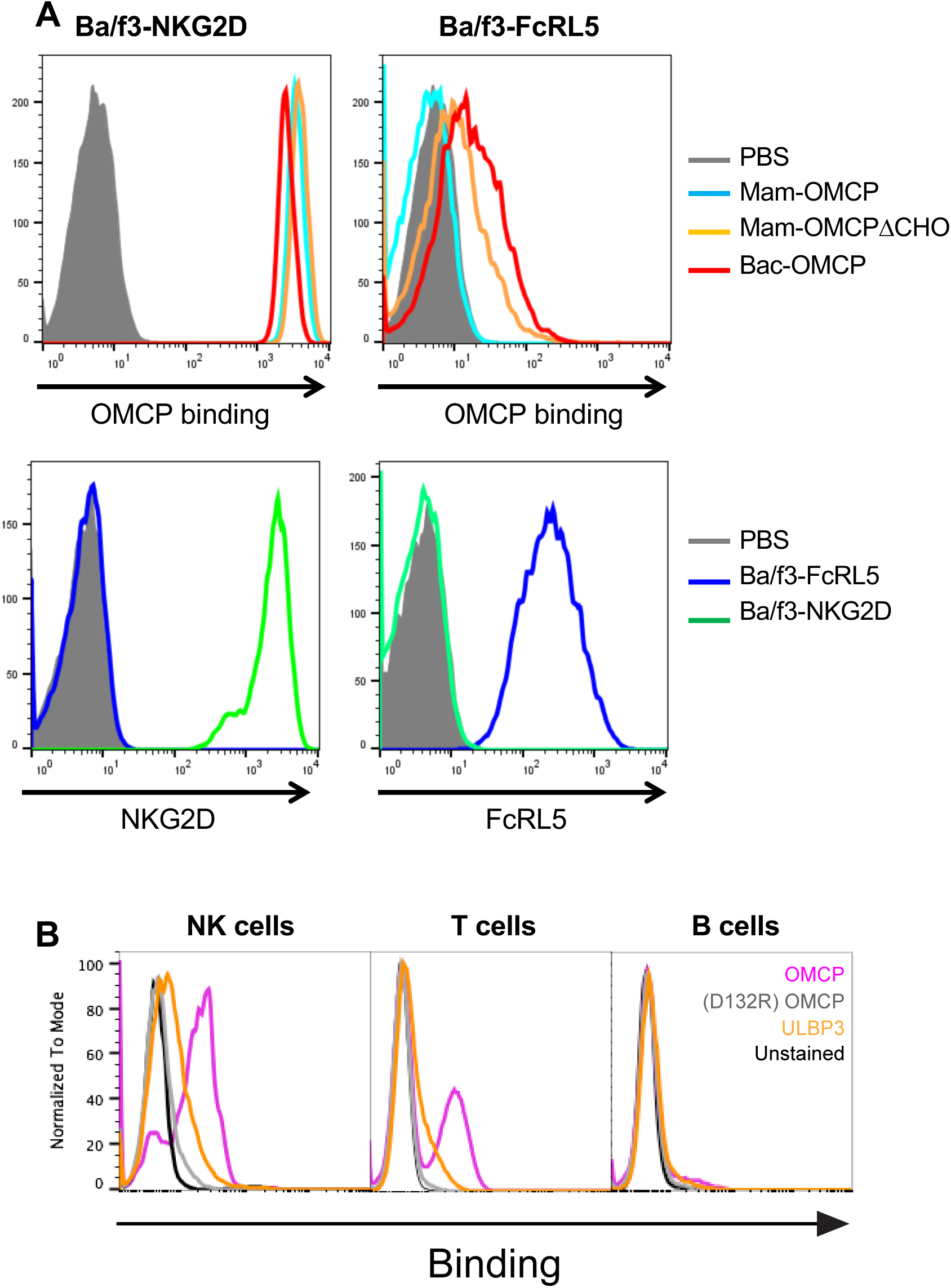
Glycosylated OMCP binds to NKG2D only. **(A)** Sorted Ba/Fc cells transduced (EGFP positive) with NKG2D or FcRL5 were stained with biotinylated monomeric OMCPs or with antibodies to NKG2D or FcRL5. A representative experiment is shown. **(B)** Pentamers of OMCP, (D132R) OMCP, or ULBP3 were incubated with ficoll-purified human PBMCs. PBMCs were stained with antibodies to CD56, CD19, CD3, and CD16. NK (CD56+, CD3-), T (CD3+), and B (CD19+, CD3-) cells were then assessed by flow cytometry for protein binding. Representative results from four independent experiments.

### Structural basis for species-specific affinity

Previously, the 14-fold higher affinity of OMCP for human vs murine NKG2D was mapped to three amino acid substitutions in the β5’-β5 loop of NKG2D, abbreviated L2 [38]. In addition to the substitutions themselves (I182V, M184I and Q185P), the position of the loop between NKG2D orthologs differs. L2 in human NKG2D is bent towards the center of the concave binding cavity compared to L2 of murine NKG2D. Superimposition of murine NKG2D onto the human NKG2D-OMCP structure reveals that the contacts between OMCP and Met184 (mNKG2D residue I200) in NKG2D^B^ and between Met184 (I200) and Glu185 (P201) in NKG2D^A^ would be altered due to the different position of the murine β5’-β5 loop (Figure 6A-B). This alteration would disrupt contacts with three residues in OMCP H2a, three residues in H2b and Arg66 within the α1 domain. Of the contact residues of L2, Met184 makes the most significant contacts in both NKG2Ds (Table 2)(Figure 6C). Critically, of the 58 NKG2D sequences available in GenBank, 54 conserve the Met184 and Glu185 found in the high affinity human NKG2D (Figure 6D).

**Figure 6.**
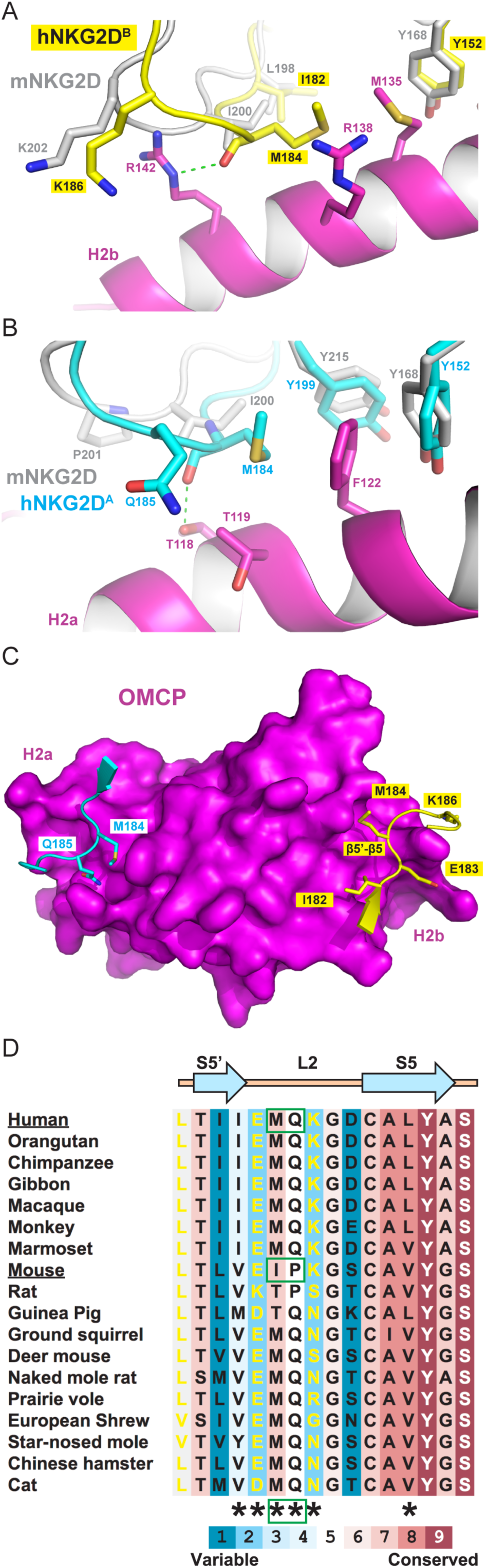
Differences in the β5’-β5 loop (L2) of human and murine NKG2D. **(A-B)** Superimposition of mNKG2D (grey) (PDB ID: 1HQ8) with the structure of OMCP-hNKG2D (yellow and cyan). Core binding residues Y152 (Y168) and Y199 (Y215) are positionally conserved, while core binding residue M184 (I200) is not. **(C)** Surface representation of OMCP (magenta) interacting with the β5’-β5 loop. **(D)** Conservation of M184 and Q185. Only the NKG2D of mice, rats, guinea pigs, and flying foxes (not shown) differ. Conservation score is as computed by the ConSurf server.

Eighteen OMCP variants have been described between different CPXV and MPXV strains [51]. In this study, we have crystallized OMCP from the Brighton Red strain of CPXV which has >60% sequence identity with the highly conserved sequence of the other 17 OMCP variants, collectively termed OMCP_mpx_. Of the 12 OMCP contact residues observed, 9 are identical to OMCP_mpx_. Of the remaining contacts, all three are conservative hydrophobic substitutions (I49L, T118I and M135I) (Figure 7). OMCP_mpx_ binds to NKG2D and the substitutions in the NKG2D contact residues are unlikely to grossly affect the affinity of OMCP_mpx_ for NKG2D [37].

**Figure 7.**
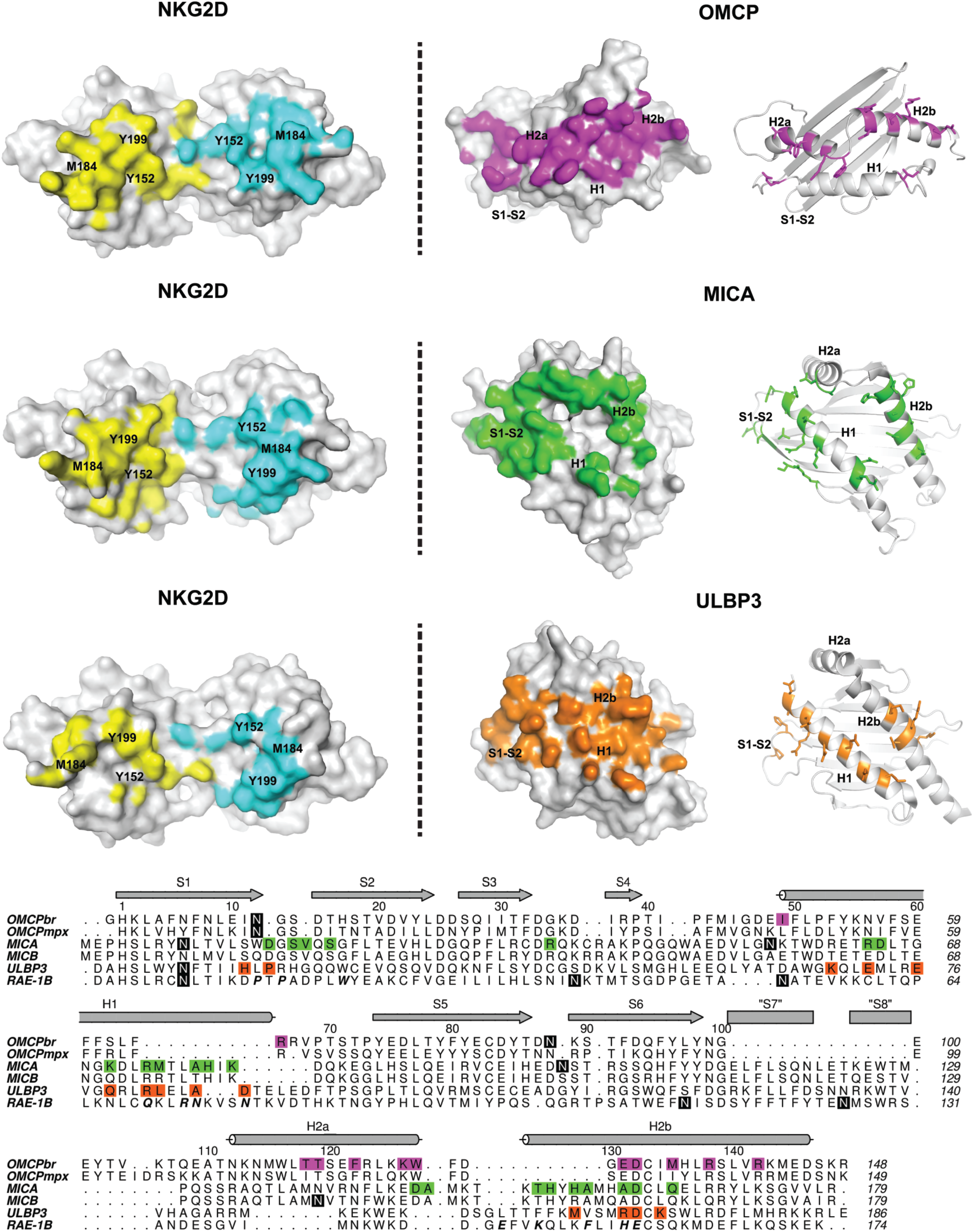
A novel NKG2D binding adaptation. Surface representation of NKG2D and surface and cartoon representations of OMCP, MICA and ULBP3. Buried surface areas for NKG2D^A^ and NKG2D^B^ are indicated in cyan and yellow, respectively. Buried surface area by NKG2D is indicated for OMCP (magenta), MICA (green), and ULBP3 (orange). The core binding residues of NKG2D and NKG2D-binding elements of NKG2DLs are indicated. Alignment by secondary structure of NKG2DLs (PDB ID: OMCP (4FFE), MICA (1HYR), MICB (1JE6), ULBP3 (1KCG) and RAE-1β (1JSK)). Contact residues are indicated for OMCP (magenta), MICA (green), ULBP3 (orange) and RAE-1β (bold and italics). Secondary structure elements are noted above the sequence (arrow for beta sheets, cylinders for alpha helices). Predicted glycan sites are highlighted in black.

### A novel NKG2D-binding adaptation

Host NKG2DLs have low sequence identity but overall similar structures, with MHCI-like platform domains binding diagonally across the symmetric binding groove created by the NKG2D homodimer [13, 42, 52]. Host ligands contact one NKG2D half site with H1 and the S1-S2 loop, and contact the second NKG2D half site with H2b. Despite the similar MHCI-like fold, OMCP binds the NKG2D binding groove in a novel orientation, rotating ∼45° relative to host NKG2DLs (Figure 7). Instead of using H1 and S1-S2 loop like host ligands, OMCP has replaced these contacts with H2a. This rotation leads to the helices of OMCP being perpendicular to the NKG2D binding groove, instead of lying diagonally across it.

Two unique rearrangements of H2a and H2b make the OMCP orientation possible. The α2 helices of OMCP and host NKG2DLs are discontinuous, with the two shorter helices hinged relative to each other. For host ligands, the angle between H2a and H2b is ∼90°, positioning H2a away from the NKG2D interface. In contrast, OMCP has increased the hinge angle between the helices by ∼20°, leading to a α2 helix that is flatter relative to the beta sheet of OMCP. The flattening of the α2 helix allows H2a and H2b to closely complement the concave binding groove of the NKG2D homodimer (Figure 2B). The tight fit of the α2 helix for NKG2D is reflected in the high shape complementarity (0.77) and buried surface area (2,612 Å^2^). In contrast, host NKG2DLs have shape complementarity ranging from 0.63-0.72 and buried surface areas ranging from 1,700-2,180 Å^2^ [44, 45, 47].

The second unique feature of the α2 helix is the separation of H2a and H2b relative to each other. This region also contains a translation that completely separates H2a and H2b into two distinct helices. This translation is critical for NKG2D binding because it allows each helix to be directly centered on the core binding sites of each NKG2D monomer (Figure 7). This creates a symmetric binding site on OMCP that recognizes the symmetric binding groove created by the NKG2D dimer. The symmetry between OMCP and NKG2D binding is in stark contrast to the canonical binding of an asymmetric host ligand to the symmetric NKG2D binding groove [52]. However, one element of asymmetry remains in the OMCP-NKG2D interaction because each NKG2D half-site recognizes an OMCP helix in a different N- to C-terminal orientation, demonstrating again the flexibility of NKG2Ds rigid adaptation recognition [42, 53].

The contact sites between NKG2D and host NKG2DLs are made up of two patches centered on the core binding sites of NKG2D and H1/S1-S2 loop and H2b of NKG2DLs [42]. As a result, the interface of NKG2D with NKG2DLs is discontinuous, particularly in the center of the NKG2D binding groove (Figure 7). Due to the unique orientation of OMCP, H2a and H2b make continuous contacts along the entire NKG2D binding groove (Figure 7). The sidechains of OMCP Lys126, Trp127, Glu131 and Asp132 make contacts with residues in the center of the NKG2D binding groove and bridge the core binding sites on each NKG2D monomer (Figure 2B). In particular, OMCP Trp127 is directed towards the center of the NKG2D dimer and makes hydrophobic contacts with residues on both NKG2D monomers, effectively closing any gaps in the binding interface.

### Signaling of NKG2D upon ligand engagement

CPXV and MPXV-infected cells secrete OMCP, which can act as an NKG2D-antagonist and block *in vitro* NKG2D-mediated NK cell killing of target cells [37]. This immune evasion strategy is reminiscent of cancer induced-NKG2DL shedding. Some cancer cells proteolytically cleave NKG2DLs from the cell surface using matrix metalloproteinases (MMPs), simultaneously preventing NKG2D-bearing lymphocytes from targeting the cancer cell, as well as creating soluble NKG2DLs to inhibit NKG2D in *trans*. Cell-associated NKG2DLs trigger NKG2D effector functions (Figure 8A), while cancer-induced, soluble NKG2DLs block NKG2D function (Figure 8B). Like shed NKG2DLs, OMCP is soluble and blocks NKG2D function in *trans* [37] (Figure 8C). Unlike host NKG2DLs, OMCP binds NKG2D with a novel orientation. We therefore asked whether OMCP could serve as a NKG2D agonist in the context of the cell membrane, analogously to host NKG2D ligands. Since OMCP is a secreted protein, an artificially cell-associated OMCP was constructed by using a heterologous GPI anchor from Thy1.1 [37] (Figure 5D). To measure NKG2D-mediated cell killing, we stably transduced Ba/F3 cells with retroviral vectors expressing either the OMCP-Thy1.1 construct or host NKG2DLs. OMCP-Thy1.1-expressing target cells were killed equivalently to host NKG2DL-transduced target cells, indicating that despite its altered binding orientation, cell-associated OMCP was able to activate NKG2D signaling (Figure 8E). Thus, OMCP must be secreted lest it active NKG2D-effector functions itself, despite potential loss of efficacy due to diffusion.

**Figure 8.**
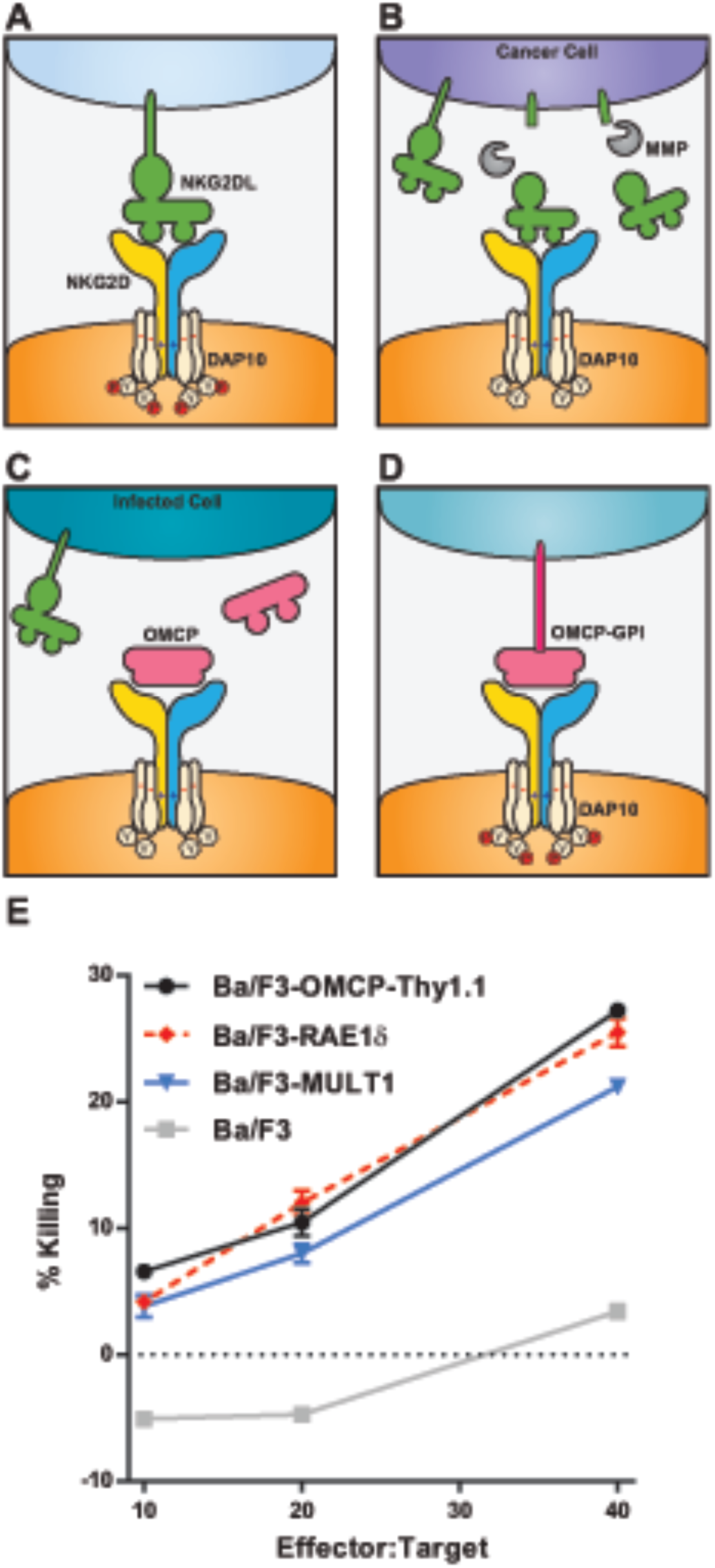
Activation of NK cells by cell-associated OMCP. Model depicting NKG2D interaction with **(A)** host, **(B)** cancer-induced, **(C)** viral, or **(D)** chimeric ligands. Binding interactions that lead to NKG2D-mediated signaling are indicated by DAP10 tyrosine phosphorylation (red filled circles). **(E)** IL-2-activated splenocytes were used as cytotoxic effectors against stably transduced Ba/F3 cell lines. Splenocytes were activated with 200 U/ml of IL-2 for 24 hours. Labeled target cells were co-incubated with activated splenocytes for 4 hours at effector:target ratios of 10:1, 20:1, and 40:1. Killing was measured by incorporation of 7AAD by CFSE-labeled target cells using flow cytometry. Representative data from five independent experiments is shown

## Discussion

Pathogens deploy multiple mechanisms to subvert the host immune response. We have previously reported that the poxvirus protein OMCP binds to two different murine immune receptors, NKG2D and FcRL5. Here, we demonstrate for the first time that OMCP facilitates cowpox virus virulence *in vivo* thorough antagonism of NKG2D. Surprisingly, we found no impact of FCRL5 *in vivo* during infections with OMCP-deficient CPXV or D132R CPVX (expressing OMCP with a mutation selectively abrogating binding with NKG2D). Investigations into the absence of a phenotype in the FCRL5-deficient mice revealed that mammalian-derived, glycosylated OMCP did not bind to FCRL5 in contrast to bacterially-expressed OMCP. Treatment of the mammalian-derived OMCP with EndoF removed the N-linked glycan and exposed a portion of OMCP that would not be present during *in vivo* infections. Surprisingly, this normally hidden surface of OMCP was able to interact with FcRL5-expressing cells. This observation has implications more broadly for the use of bacterially-expressed proteins in receptor-ligand studies. However, these data highlight the importance of OMCP in subverting NKG2D-mediated control of cowpox virus infection. In the absence of OMCP antagonism of NKG2D, cowpox virus was significantly attenuated. Cowpox virus inhibits the cell surface expression of MHCI to avoid recognition by effector T cells[30, 31]. The loss of MHCI expression on the cell surface induces “missing self” and enhanced recognition by NK cells. In the absence of OMCP antagonism of the activating NK cell receptor NKG2D, infected cells become more susceptible to NK cell-mediated killing.

While many viruses have adopted a general mechanism of NKG2D-sabotage by trying to retain multiple host-encoded NKG2D ligands within the infected cell, CPXV and MPXV take the very different approach of targeting NKG2D directly. Since NKG2D is monomorphic, this mechanism has the significant advantage of requiring a single protein to prevent NKG2D recognition of the infected cell. The large number of sequence-divergent host NKG2DLs and their associated polymorphisms are thought to be driven by selection from pathogen-encoded NKG2DL antagonists [14]. Likewise, viral NKG2L antagonists are under selective pressure from the diverse host NKG2DLs in a continual cycle of adaptation. Due to the need to recognize multiple NKG2DLs, NKG2D has a limited mutational space to adapt. The limited ability of NKG2D to mutate is yet another advantage of OMCP directly targeting NKG2D, instead of NKG2DLs.

Similarly to OMCP, some cancer cells shed host NKG2DLs to create their own soluble NKG2D antagonists. However, this strategy has the additional benefit of removing host NKG2DL from the surface of cancer cells. In contrast, CPXV and MPXV lack a known mechanism of blocking host NKG2DL surface expression. Secreted OMCP must then be able to compete efficiently against the high local concentration of multiple host NKG2DLs on the infected cell, as well as against diffusion away from the infected cell. One possible way to increase OMCP’s ability to compete with host ligands would be to increase the avidity of OMCP by having multiple NKG2D-binding domains. However, a multimeric OMCP could crosslink NKG2D and potentially trigger NKG2D-mediated killing. Therefore, secreted OMCP must be monomeric to prevent aberrant NKG2D signaling. Thus to compensate for these deficiencies, OMCP must have the highest possible affinity to effectively compete against cell-associated host NKG2DLs [37, 38]. The half-life of ligand-receptor interactions correlate well with physiological competitiveness [54]. OMCP binds human and murine NKG2D with half-lives of 348 and 54 seconds, respectively, compared to half-lives of 1.5-18 seconds for most NKG2DLs [38, 45, 55]. Indeed, the increased half-life for NKG2D allows OMCP to effectively antagonize NKG2D-mediated immunity in our murine infection models despite the lower affinity of the murine NKG2D receptor for OMCP compared to human NKG2D receptor. CPXV infections in primates or rodents that express the higher affinity NKG2D receptor would likely be protected from NKG2D-mediated immunity by OMCP to a greater degree.

To understand the molecular basis for the long half-life of OMCP for NKG2D, we previously determined the structure of OMCP alone, and here, we report the structure of OMCP bound to NKG2D. The structure of OMCP alone was grossly similar to that of host NKG2D ligands, containing an atypical MHCI-like platform domain. Host NKG2D ligands bind with the helices of their platform domains oriented diagonally within the symmetric binding groove of NKG2D. Thus it was expected that OMCP was a viral mimic of host NKG2D ligands and would interact with NKG2D analogously.

The structure of OMCP-NKG2D instead revealed a novel orientation for an NKG2D ligand in the NKG2D binding groove. Alterations within the α2 domain helix allow OMCP to arrange its helices perpendicularly within the binding groove. This reorientation places the H2a and H2b helices directly in contact with the core binding sites of NKG2D and also forms the largest and most continuous binding interface with NKG2D. Because the forces (hydrogen bonds, van der Waals, hydrophobic interactions) that mediate protein-protein interactions are individually weak, a large, continuous interface with high shape complementary allows for a cumulatively strong interaction between proteins. This change in the binding orientation of OMCP reveals how the MHCI-like platform used by host ligands can be adapted by a pathogen to enhance NKG2D binding.

Since host NKG2DLs and OMCP have a similar MHCI-like platform, it is reasonable to wonder why no host ligand has evolved an analogous high-affinity interaction with NKG2D. One likely reason is that the host immune response must be carefully calibrated to balance the need for protection against the threat of autoimmunity. Since the expression of NKG2DLs on the cell surface signals for effector functions, even a small amount of high affinity host ligand on the cell surface could trigger an immune response, and the resulting tissue damage could be deleterious for the host. Indeed, NKG2D-expressing cells and/or aberrant expression of host NKG2DLs have been implicated in diabetes, celiac disease and rheumatoid arthritis [56-59]. Viruses are not constrained by autoimmune selective pressures. Therefore, CPXV and MPXV were free to evolve a viral NKG2DL with the highest possible affinity to maximize immune evasion potential.

Interestingly, OMCP triggers NKG2D signaling when attached to a target cell membrane, despite the novel orientation of OMCP relative to host NKG2DLs. The interaction of host NKG2DLs with the dimeric NKG2D bears broad structural similarity to the interaction between MHC molecules with their cognate T cell receptors (TCRs). In both cases, the NKG2DL/MHC lies diagonally across the surface created by the dimeric NKG2D/TCR. However, there are several examples of MHC-TCR complexes that, like OMCP-NKG2D, interact with unconventional orientations [60-64]. Several of these complexes involved autoimmune MHC-TCR complexes that were tilted or rotated outside of the normal range for MHC-TCR complexes [60, 64]. While these receptors could induce TCR signaling at high MHC concentrations, they failed to assemble characteristic immunological synapses [65]. A striking example of unconventional binding was found when an *in vitro* peptide library-MHC-TCR (H2-L^d^-42F3) screen produced a p3A1-H2-L^d^-42F3 complex with an interface rotated ∼40° relative to other H2-L^d^-42F3 complexes. This rotation places the TCR nearly parallel with the MHC peptide-binding groove and shifted the interface center almost entirely on one of the MHC α helices - an orientation strikingly similar to the interface of OMCP-NKG2D [64]. Interestingly, the p3A1-H2-L^d^-42F3 complex failed to induce TCR signaling [64]. Thus, unlike OMCP/NKG2D, the orientation of MHC relative to TCR is an important factor for signaling.

OMCP-NKG2D and p3A1-H2-L^d^-42F3 have opposite signaling outcomes, despite having very similar orientations. TCR signaling requires co-receptor binding to either the α2/β2 or α3 domains of MHCII or MHCI, respectively. The failure of p3A1-H2-L^d^-42F3 to signal, and of other unconventional MHC-TCR complexes to form true immunological synapses, is potentially due to the inability of co-receptors to form correct quaternary structures for signaling [63, 64, 66]. Signaling by NKG2D is not known to require co-receptor stimulation and the majority of NKG2DLs lack the co-receptor binding α2/β2 or α3 domains of true MHC molecules. This difference in co-receptor dependency likely explains why OMCP (when attached via GPI anchor) is still competent to stimulate NKG2D-signaling compared to MHC-TCR complexes with unconventional binding orientations. Further, it suggests that clustering of NKG2D on the cell surface is the major determinant of NKG2D-mediated activation.

## Materials and Methods

### Cell lines and mice

The generation of Ba/F3 transductants expressing NKG2D, FcRL5, RAE1δ, MULT1, and OMCP-Thy1.1 was previously described [37, 67]. C57BL/6 (B6) mice were obtained from the National Cancer Institute (Charles River, MA). NKG2D-deficient mice on a B6 background were a kind gift from Bojan Polic (University of Rijeka, Croatia) [68], and FcRL5-deficient mice on a B6 background were obtained from Riccardo Dalla Favera (Columbia University). Mice were maintained under specific pathogen–free conditions and used between 8 and 12 weeks of age. Female mice utilized in survival studies were observed daily after infection for 28 days and moribund mice were euthanized per institutional guidelines.

### Ethics Statement

All experiments were conducted in accordance with institutional guidelines for animal care and use based on the Guide for the Care and Use of Laboratory Animals of the National Institutes of Health. The protocol was approved by the Animal Studies Committee at Washington University (#20110104). Human PBMC were collected under IRB approved protocol #201110275 at Washington University in St Louis. Informed consent was obtained from all volunteers donating blood.

### CPXV and infection of mice

Brighton Red strain CPXV was obtained from the ATCC and propagated in BS-C-1 cells. Virus was purified from infected BS-C-1 cell lysates by centrifugation through a 36% sucrose gradient. The titer of the viral stock was determined using a standard plaque assay on BS-C-1 cells [69, 70]. Intraperitoneal (i.p.) infections were done at a dose of 2×10^6^ pfu CPXV/mouse unless otherwise indicated. For intranasal infections, mice were first anesthetized i.p. with ketamine/xylazine short acting sedative. Mice were infected with the appropriate innoculum of virus in a 50µL volume intranasally. All i.n. infections were done at a dose of 2.5×10^4^ pfu CPXV/mouse unless otherwise indicated.

### Generation of recombinant cowpox viruses

Recombinant virus lacking expression of OMCP (ΔV018) and its revertant were produced by transient dominant selection [71, 72]. A PCR product encoding the VACV p7.5 5’ UTR and promoter was amplified and spliced to another PCR product encoding the *E. coli gpt* sequence. The final product was cloned into plasmid pUC19 to create pUC19 p7.5 *gpt*. OMCP was then amplified in two fragments in order to insert two tandem early termination codons. This product was cloned into pUC19 7.5 *gpt* to create pUC19 p7.5 *gpt* OMCPSTOP. The revertant plasmid pUC19 p7.5 *gpt* OMCP was generated by excising the OMCPSTOP sequence and replacing it with wild-type OMCP. OMCP D132R mutant CPXV was generated in a similar manner with primers to insert a point mutation at D132. The mutated D132R construct was also spliced to *E. coli gpt* and cloned for transient dominant selection.

Confluent CV-1 cells were then infected with CPXV (for CPXV ΔV018 or D132R generation) or CPXV ΔV018 (for revertant generation) at an MOI of 0.5. Two hours post-infection, the cells were transfected with pUC19 p7.5 *gpt* OMCPSTOP for CPXV ΔV018, pUC19 p7.5 *gpt* OMCP-D132R for D132R generation, or pUC19 p7.5 *gpt* OMCP for revertant generation using Lipofectamine2000. The infected/transfected cells were lysed after 48 hours, and *gpt*+ viruses were selected by infecting confluent B-SC-1 cells with the transfected virus stock in *gpt* selection medium (DMEM, 2% FCS, 25 μg/mL mycophenolic acid, 250 μg/mL xanthine, and 15 μg/mL hypoxanthine). *Gpt*+ viral plaques underwent two further rounds of selection in *gpt* selection media and two subsequent rounds of purification in non-selective media. Finally, plaques were tested for their ability to grow in both selective and non-selective media. Plaques that grew only in non-selective media (indicating loss of the *gpt* marker) were picked and underwent a total of five plaque purifications. Virus stocks were subsequently propagated in B-SC-1 cells and titered following standard protocols using B-SC-1 cells [70].

### Identification of NKG2D-binding null mutant D132R

A high throughput *in vitro* selection approach based on combinatorial cell surface display was utilized to identify NKG2D-binding null mutants. The sequence of OMCP was globally mutagenized using error-prone PCR, and the mutated amplicons were spliced to a signal-less Thy1.1 cDNA via overlap extension PCR. This library of mutated OMCPs fused to unmutated Thy1.1 was cloned into the pMXs-IRES-EGFP retroviral transfer vector (kind gift of Toshio Kitamura, University of Tokyo) to generate a molecular library for transduction into Ba/F3 cells. The transductants were then sorted for green fluorescence and anti-Thy1.1 expression to yield a cellular library whose members all had surface expression of OMCP, filtering out mutations giving frameshifts, premature stop codons, and folding-incompetent OMCP. This OMCP library was sorted for NKG2D binding using NKG2D-tetramers. Sorted cells were cloned by limiting dilution and analyzed. The retroviral cassettes of cells lacking or having reduced NKG2D-binding activity were amplified and sequenced. Utilizing this approach, we identified Asp132 as a critical residue for NKG2D binding.

### Protein expression and purification

OMCP_BR_ and human NKG2D expression constructs were previously described [38]. The (D132R) OMCP_BR_ protein was prepared identically to WT OMCP_BR_. (23D/95D) OMCP-NKG2D complex was reconstituted by oxidative co-refolding from purified inclusion bodies, as described previously [38]. Refolded protein was slowly diluted 10-fold with water and captured on a 5 ml HiTrap Q HP column (GE Healthcare) using a Profinia instrument (Bio-Rad). The captured protein was washed with 50 mM Tris, pH 8.5, 20 mM NaCl and bulk eluted with 50 mM Tris, pH 8.5, 250 mM NaCl. The eluted protein was then concentrated and further purified by gel filtration chromatography on a Superdex S75 column (16/60; Amersham Biosciences). Fractions containing mono-dispersed OMCP-NKG2D complex (∼50 KDa) were pooled and buffer exchanged into 25mM Ammonium acetate pH 7.4.

Mammalian-derived proteins were expressed by transient transfection of HEK293F cells (Life Technologies) using PEI. The culture medium was collected 3 days and 6 days after transfection, and then purified by standard Ni-NTA chromatography in accordance with the manufacturer’s protocol (Gold Biotechnology) and were subsequently buffer exchanged into phosphate-buffered saline (PBS). Pentamerized proteins were made by fusing the target protein with a C-terminal cartilage oligomeric matrix protein (COMP) domain [73].

### Crystallization, data collection and processing

Native protein crystals were grown by hanging drop vapor diffusion at 20°C by streak seeding into a well solution containing 15% PEG 3350, 0.2M MgCl_2_, 0.1M Bis-Tris pH 6.75. Crystals were cryoprotected with well solution containing 15% glycerol before flash freezing directly in a liquid nitrogen bath. Diffraction data were collected at the Advanced Light Source synchrotron (beamline 4.2.2). Native (23D/95D) OMCP-hNKG2D crystal diffraction data were collected at 100 K and at a wavelength of 1.00004 Å. Additional diffraction data statistics are summarized in Table 1. Data processing with HKL2000 [74] showed the crystals belonged to the primitive monoclinic space group P2_1_ (space group #4). The asymmetric unit of the crystal contained two copies of the (23D/95D) OMCP-hNKG2D complex.

### Model building and refinement

The structures of human NKG2D (1MPU)[49] and OMCP (4FFE)[38] were used as search models for molecular replacement through Phenix [75]. Reiterative refinement and manual rebuilding were performed using Phenix and Coot [76], respectively. Both 2Fo-Fc and Fo-Fc maps were used for manual building and to place solvent molecules. The final model yielded an R_work_ of 16.6% and R_free_ of 21.4%, with 4% of all reflections set aside for free R factor cross-validation. Progress in refinement was also measured using the MOLPROBITY webserver [77]. The final Ramachandran statistics for the model were 98% favored and 0% outliers. Additional refinement statistics are summarized in Table 1. Images of structures were produced using the program PyMol [78].

### Structure analysis

Analysis of the contact residues, buried surface area and shape complementarity of the OMCP-NKG2D interface were carried out using the programs Ligplot+ [79], PISA [80] and SC [81]. Structural programs as compiled by the SBGrid consortium [82]. Analysis of NKG2D conservation was performed using the ConSurf server [83-86]. GenBank numbers for species used in Consurf analysis are: Humans (30749494), Borean orangutan (21902299), Chimpanzee (57113989), Gibbon (332232684), Macaque (355785888), Green Monkey (635063485), Common marmoset (380848799), Mouse (148667521), Brown rat (149049263), Guinea Pig (348569092), Ground squirrel (532114387), Deer mouse (589967905), Naked mole rat (512868733), Prairie vole (532053033), European Shrew (505834608), Star-nosed mole (507978716), Chinese hamster (537136230), and Cat (410963826).

### Atomic coordinates

The atomic coordinates (accession code 4PDC) have been deposited in the Protein Data Bank, Research Collaboratory for Structural Bioinformatics (Rutgers University, New Brunswick, NJ)

### *In vitro* NK cell killing assays

Splenocytes from C57BL/6 mice were preactivated with 200 U/ml IL-2 for 24 hours and used as cytotoxic effectors against stably transduced Ba/F3 cell lines in standard killing assays. Target cells were carboxyfluorescein succinimidyl ester (CFSE) labeled and co-incubated with activated splenocytes at 37°C, 5% CO2 for 4 hours at effector:target ratios of 10:1, 20:1, and 40:1. Killing percentage was determined by incorporation of the dead cell exclusion dye 7-amino-actinomycin D (7AAD) in the CFSE+ target population as assessed by flow cytometry. Percent specific lysis was calculated using the formula [(experimental dead % - background dead %) / (maximum release dead % - background dead %)] x 100. Single cell suspensions of splenocytes used in killing assays were generated using standard protocols[87].

### Antibodies, and Flow cytometry

Recombinant OMCP, D132R OMCP, and West Nile Virus glycoprotein domain III (DIII) were produced and biotinylated as previously reported [37, 88]. APC labeled tetramers were made by incubating biotinylated OMCP protein with APC-streptavidin in a 4:1 molar ratio for 10 min at room temperature. APC eFlour 780 anti-CD3 (145-2C11) and eFlour 450 anti-CD19 (1D3) were purchased from eBioscience (San Diego, CA). FITC CD19 (1D3) and PE anti-NK1.1 (PK136 were from BD Biosciences (San Jose, CA). FITC anti-CD5 (53-7.3) was from Biolegend (San Diego, CA). Antibodies and staining of human PBMCs were performed as previously reported [39]. Single cell suspensions of splenocytes and isolation of peritoneal cells and Ficoll purification of human PBMCs were performed using standard protocols [87, 89]. To block nonspecific binding of antibodies to FcRs, murine cells were incubated in 2.4G2 (anti-FcγRII/III) supernatants (hybridoma from ATCC) prior to staining with labeled Abs. Data was collected on a FACScan flow cytometer (BD Pharmingen) with DxP multicolor upgrade (Cytek Development, Fremont, CA) and FlowJo CE software (TreeStar, Ashland, OR) or FACSCalibur flow cytometer (BD Biosciences) with CellQuest collection software (BD Biosciences). Data analysis was performed using FlowJo software (TreeStar).

### Plaque assay

Tissues harvested from mice were weighed and collected in tubes containing 1mL of DMEM and stored at −80°C until processing. Frozen organs were homogenized using glass dounce homogenizers and collected in 1mL of DMEM. The tissue homogenate was serially diluted in D2.5 media (DMEM, 2.5% FBS, 1% L-glutamine, 1% penicillin-streptomycin, 1% non-essential amino acids) and used in standard plaque assays on BS-C-1 cells [69]. Each organ was titered on 5×10^5^ BS-C-1 cells plated in wells of 6 well plates in duplicate. Infected cell cultures were maintained in D2.5 media at 37°C, 5% CO_2_ for 48 hours. Cells were stained at 48 hours post infection using a crystal violet solution and plaques were counted. Viral titers were calculated from a minimum of 3 dilutions done in duplicate.

### Statistical analysis

Data analysis was done with Microsoft Excel and GraphPad Prism (GraphPad Software, La Jolla, CA). Unless otherwise noted, unpaired, two-tailed t tests were used to determine statistically significant differences. Error bars in the figures represent SDs from the mean value. The Log-rank (Mantel-Cox) test was used in the comparison of all Kaplan Meier survival curves.

## Acknowledgements

E.L., X.W., C.N., and D.H.F. were supported in part by NIH/NIAID R01 AI019687, U19 AI109948 and NIAID contracts HHSN272200700058C & HHSN272201200026C. M.M.S, T.L.G, D.L, and A.R.F were supported in part by NIH/NIAID R01AI073552. A.S.K. supported by NIH PO1 AI116501. We thank Helen M. Lazear for help with illustrations

## Figure Legends

**Figure S1.**
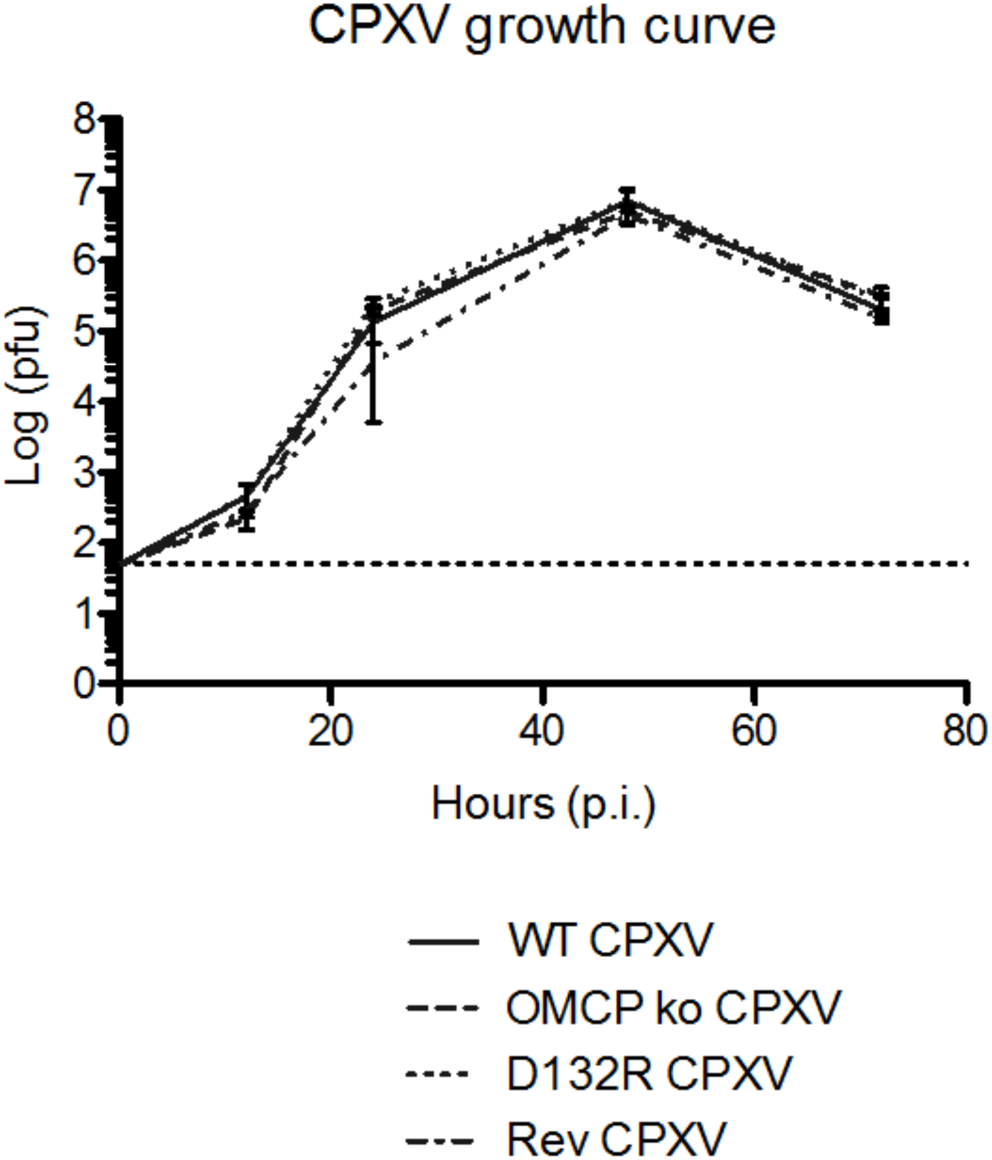
ΔV018 virus and D132R virus do not have a growth defect *in vitro*. Multi-step growth curves for WT CPXV, ΔV018, D132R, and revertant viruses show that all four cowpox virus strains have similar replication kinetics in a multi-step growth curve. B-SC-1 cells were infected at an MOI of 0.01 with the indicated viruses. Infected cells were harvested at the indicated time points and freeze-thawed three times. Lysates were titered by plaque assay on B-SC-1 monolayers. Results are representative of three experiments done in duplicate.

**Figure S2.**
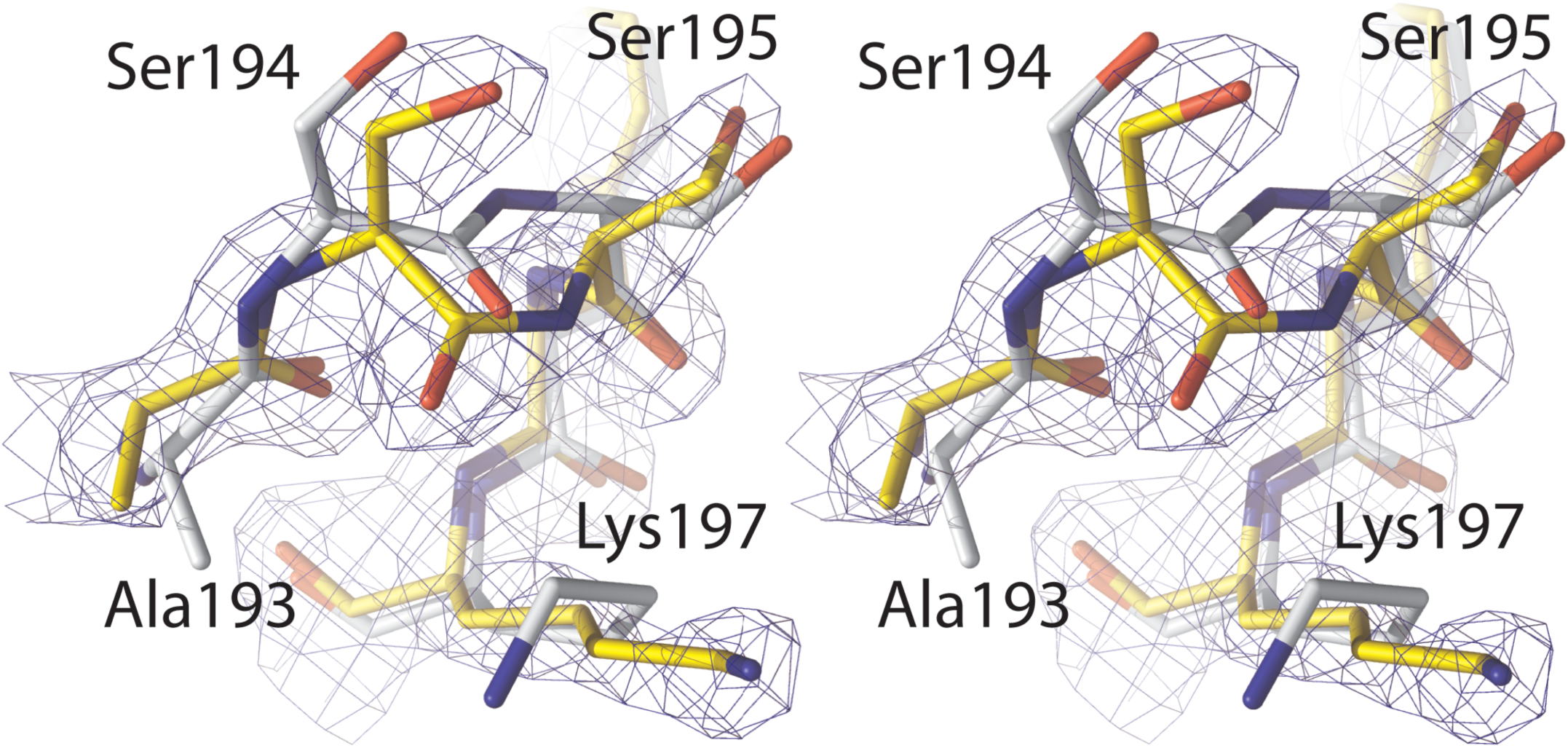
Electron density supporting a *cis* peptide conformation. Stereo view of the β5-β6 loop of hNKG2D. Residues 193-Ala-Ser-Ser-Phe-Lys-197 is displayed for the OMCP-hNKG2D structure (yellow) and the structure of hNKG2D alone (grey). The 2Fo-Fc map for OMCP-hNKG2D is displayed at 2σ. *Cis* conformations are underrepresented in crystal structures and are often under reported due to resolution limits and assumptions made during structure refinement [90, 91].

**Figure S3.**
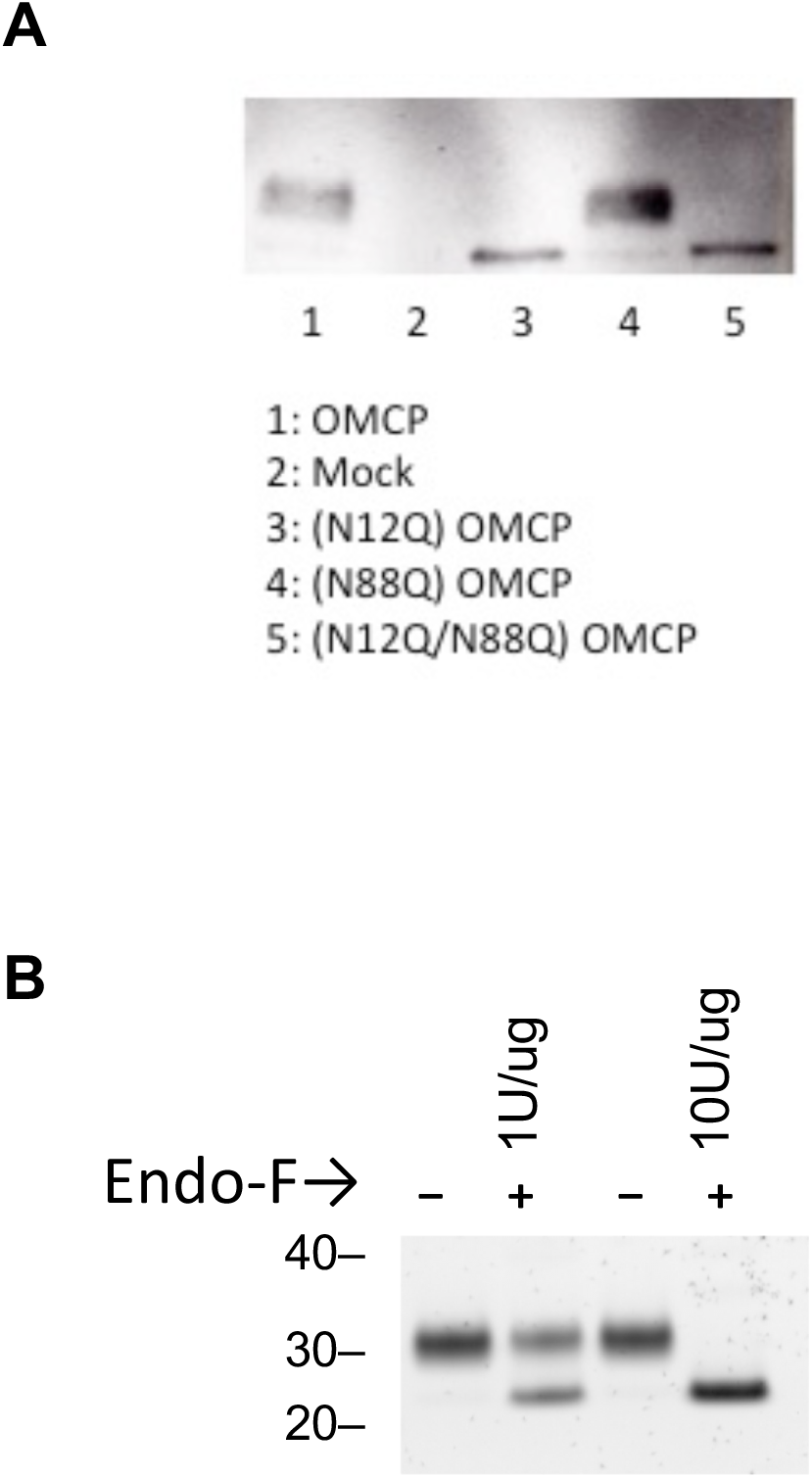
Glycosylation of mammalian-expressed OMCP. OMCP has two putative N-linked glycosylation sites, N12 and N88 **(A)** WT, glycosylated OMCP migrates slowly as a diffuse band. OMCP mutated at either N12 or N12/N88 migrate quickly as sharp bands. Mutation of N88 alone does not effect OMCP migration. **(B)** N-linked glycans were completely removed from 1 ug of OMCP by incubation with 10U Endo-F in 50mM sodium phosphate (pH7.5) at 37C for 16h.

